# Shrimp endogenous viral elements (EVE) correlate with survival in white spot syndrome virus (WSSV) challenges

**DOI:** 10.64898/2026.02.28.708708

**Authors:** Suparat Taengchaiyaphum, Phasini Buathongkam, Jiraporn Srisala, Prapatsorn Wongkhaluang, Thanaporn Wongpim, Suwanchai Phomklad, Kaidtisak Kaewlok, Jaroon Inkaew, Sorawit Powtongsook, Timothy W. Flegel, Ornchuma Itsathitphaisarn, Kallaya Sritunyalucksana

## Abstract

Shrimp and other arthropods are capable of specific, adaptive immune responses to viruses based on viral copy DNA (vcDNA) fragments in the host genome called endogenous viral elements (EVE). These may produce negative sense RNA transcripts leading to an RNA interference (RNAi) defense response against cognate viruses. We first reported high-frequency-read sequences (HFRS) of white spot syndrome virus EVE (named WSSV-EVE 4,6,8) in a WSSV-free breeding stock of whiteleg shrimp (*Penaeus vannamei*). Here we describe screening for the same HFRS-EVE in a captured giant tiger shrimp (*Penaeus monodon*) breeding stock, also free of WSSV. WSSV-EVE 4,6,8 was detected in some of the *P. monodon* stock individuals with positive or negative RNA expression. Eight broodstock individuals were selected for mating in 4 crosses. The offspring from these crosses were grown sufficiently to allow tagging and pleopod sampling for DNA and RNA analysis prior to challenge with WSSV. This allowed for Mendelian analysis of EVE inheritance and for its expression or not in the offspring, together with analysis of their relationships to survival and WSSV infection level after challenge. The results revealed that EVE inheritance was Mendelian, but that their RNA expression or not was independently controlled. In Crosses 1 and 2, all the offspring died and none of them carried 2 or more of the expressed EVE in their parental shrimp. In contrast, 100% of 10 arbitrarily selected surviving shrimp from Cross 3 and 90% from Cross 4 carried and expressed 2 or more of the 3 expressed EVE transmitted from the parental shrimp. These results reveal a potential protocol for development of viral tolerant shrimp stocks.

## 1. INTRODUCTION

There are recent reviews on the occurrence of specific adaptive immunity against both RNA and DNA viral pathogens mediated by DNA and RNA in insects (Bonning Saleh, 2021) and crustaceans (Flegel, 2022). The process has been called viral accommodation (Alday-Sanz, et al., 2020; Flegel, 2009; Flegel Pasharawipas, 1998) because it often leads to low-level persistent viral infections in survivors without signs of disease. A key feature of this defense mechanism is the development of endogenous viral elements (EVE) in the host genome, some of which may lead to host RNA interference (RNAi) responses against the cognate virus (Suzuki, et al., 2020). An additional advantage is that they can provide heritable immunity if they occur in the DNA of germ cells. Mendelian inheritance of EVE was first reported for infectious hypodermal and hematopoietic necrosis virus (IHHNV) in *P. vannamei* (Brock, et al., 2013) and later for WSSV in *P. monodon* (Taengchaiyaphum, et al., 2019; Utari, et al., 2017). These publications contained warnings that existence of EVE meant that PCR testing alone could not be used to confirm viral infections, an issue that was subsequently reviewed in detail (Alday-Sanz, et al., 2020; Saksmerprome, et al., 2021; Sritunyalucksana et al. 2025).

It was discovered that viral accommodation mechanisms included a pathway for the formation of constructs of linear viral copy DNA (lvcDNA) and circular viral copy DNA (cvcDNA) via host derived reverse transcriptase interaction with viral messenger RNA (Alday-Sanz, et al., 2020; Flegel, 2022). This provided a means of selectively isolating cvcDNA for detection of potential EVE. Upon using this process with the shrimp DNA virus IHHNV, it was discovered that EVE related to IHHNV could themselves, also give rise to cvcDNA transcripts. It, in turn, provided a method to screen for EVE by using next-generation sequencing (NGS) with cvcDNA extracts from uninfected shrimp (Taengchaiyaphum, et al., 2021). This approach was used with a breeding stock of *P. vannamei* (Taengchaiyaphum, et al., 2024) free of WSSV but derived from previous survivors of exposure to extant types of pathogenic WSSV. The resulting raw-read fragments (approximately 150 bp) with high sequence identity to extant types of WSSV were mapped intermittently across the whole WSSV genome. However, a very small portion of approximately 1,400 bp in the 300,000 WSSV genome (0.5%) exhibited a vast majority of the raw reads. We called these high frequency read sequence (HFRS) genome regions and hypothesized that they may have arisen by natural selection due to provision of tolerance to white spot disease (WSD) (Taengchaiyaphum, et al., 2024).

Here, we describe the same EVE identification process with captured broodstock of giant tiger shrimp and reveal that they exhibited a similar HFRS region to that in *P. vannamei*, supporting the proposal for natural selection of this region in two shrimp species. Further, we describe experiments with the *P. monodon* breeding stock to test our hypothesis that presence of HFRS-EVE sequences in shrimp can protect them against death from WSD. This was accomplished first by screening for individual male and female broodstock specimens that did or did not carry the WSSV-HFRS EVE we call EVE-4, -6 and -8 that matched sequences in the WSSV-HFRS genome region. This was followed by selecting broodstock carrying EVE-4,6,8 or not and expressing them or not to produce offspring that could be sampled for EVE-4,6,8 presence and RNA expression prior to WSSV challenge tests. This allowed for: (1) Mendelian analysis of EVE transmission and its RNA expression to offspring, and (2) assessment of the relationship between these factors and offspring survival. Based on our hypothesis, it was predicted that presence and expression of these EVE would be positively related to survival, despite WSSV infection.

## 2. MATERIALS AND METHODS

### 2.1. Shrimp preparation and overall experimental plan

A mating program and artificial insemination of *Penaeus monodon* broodstock captured from the wild is operated by the Shrimp Genetic Improvement Center (SGIC), Surat Thani, Thailand. Adult shrimp were screened in quarantine to select individuals that tested negative for major shrimp pathogens including WSSV. Selected adults were tagged and moved to the secure breeding center. Two pleopods were removed from each individual adult and subjected to DNA extraction. Then, a portion of the DNA from each was pooled and subjected to isolation of circular viral copy DNA (cvcDNA) that was then sent for next generation sequencing to identify endogenous viral elements (EVE) of white spot syndrome virus (WSSV), as previously described for a breeding stock of the whiteleg shrimp *Penaeus vannamei* (Taengchaiyaphum et. al., 2024).

Based on their WSSV-EVE profiles, 4 pairs of male and female specimens were selected and mated (4 crosses) to obtain 4 populations of juvenile offspring. When these juveniles reached a size of 2–3-g, 40 from each cross were arbitrarily selected and individually tagged for WSSV challenge experiments with 2 replicates of 20 shrimp (total 40) from each cross. Sex determination was not possible but was determined by subsequent genetic analysis (see below). The shrimp were then divided into 4 groups of 40 each (total 160) and housed in individual plastic cages within 500-L tanks. They were given 2-days for acclimatization before 2 pleopods were removed from each specimen and numbered corresponding to the tag on the source specimen. The pleopods were used for DNA and RNA extraction for sex determination and for WSSV-EVE profile of each specimen by PCR and RT-PCR analysis. This analysis allowed for comparison of the relationship between sex, WSSV-EVE and its transcript status of surviving and dead shrimp in the subsequent WSSV-challenges. Finally, the 4 sets were subjected to a 14-day challenges with virulent WSSV inoculum. This was followed by additional pleopod removal for individual WSSV qPCR analysis of each dead and surviving specimen selected. Moribund specimens were counted as dead specimens.

### 2.2. Origin and management of broodstock and offspring

Broodstock maintenance, mating design, artificial insemination, and offspring rearing were carried out at the Shrimp Genetic Improvement Center (SGIC), Chaiya, Surat Thani, Thailand. All procedures were performed under a strict biosecurity control system. A total of 56 broodstock individuals (25 females and 31 males), aged 11–12 months were tested for the presence of EVE 4, 6, 8 and their expression by PCR and RT-PCR. The pleopod tissue of broodstock was excised for genomic DNA and RNA extraction to facilitate screening for EVEs. Pre-screening for the absence of major pathogens was conducted using PCR/RT-PCR in accordance with the relevant World Organization for Animal Health (WOAH) reference protocols. A total of eight pathogens were tested: White Spot Syndrome Virus (WSSV), Yellow Head Virus genotypes 1–8 (YHV1–YHV8), Taura Syndrome Virus (TSV), Infectious Myonecrosis Virus (IMNV), Acute Hepatopancreatic Necrosis Disease (AHPND), *Enterocytozoon hepatopenaei* (EHP), Infectious Hypodermal and Hematopoietic Necrosis Virus (IHHNV), and Decapod Iridescent Virus 1 (DIV1).

Shrimp mating and offspring preparation steps (Fig. 1) were as follows: **(1)** *Broodstock culture conditioning*: Male and female broodstock were cultured in 30 ppt sea water, 28-30 °C for 3–4 weeks (duration depends on their readiness) using formulated broodstock feed and fresh feed (e.g., sandworms). This enhances gonadal development beyond normal rearing conditions. During this period, some female broodstock begin ovarian development, and males show increased spermatophore intensity. **(2)** *Artificial Insemination*: Once conditioning was completed and a female molted, spermatophores were extracted from the males and injected into the thelycum of the females using artificial insemination techniques. **(3)** *Eyestalk Ablation*: Three to four days after artificial insemination, eyestalk ablation was performed on the females to stimulate ovarian maturation. **(4)** *Ovarian Assessment and Spawning*: Two days post-eyestalk ablation, the females’ ovarian development was assessed. Females with stage 3 or 4 ovaries were selected for spawning, and their individual spawning cycles were monitored. **(5)** *Juvenile Culture*: Larvae were reared from the nauplius stage to juveniles weighing approximately 2–3 grams over a period of 2–3 months, before they were subjected to viral challenge tests.

**Figure 1.**
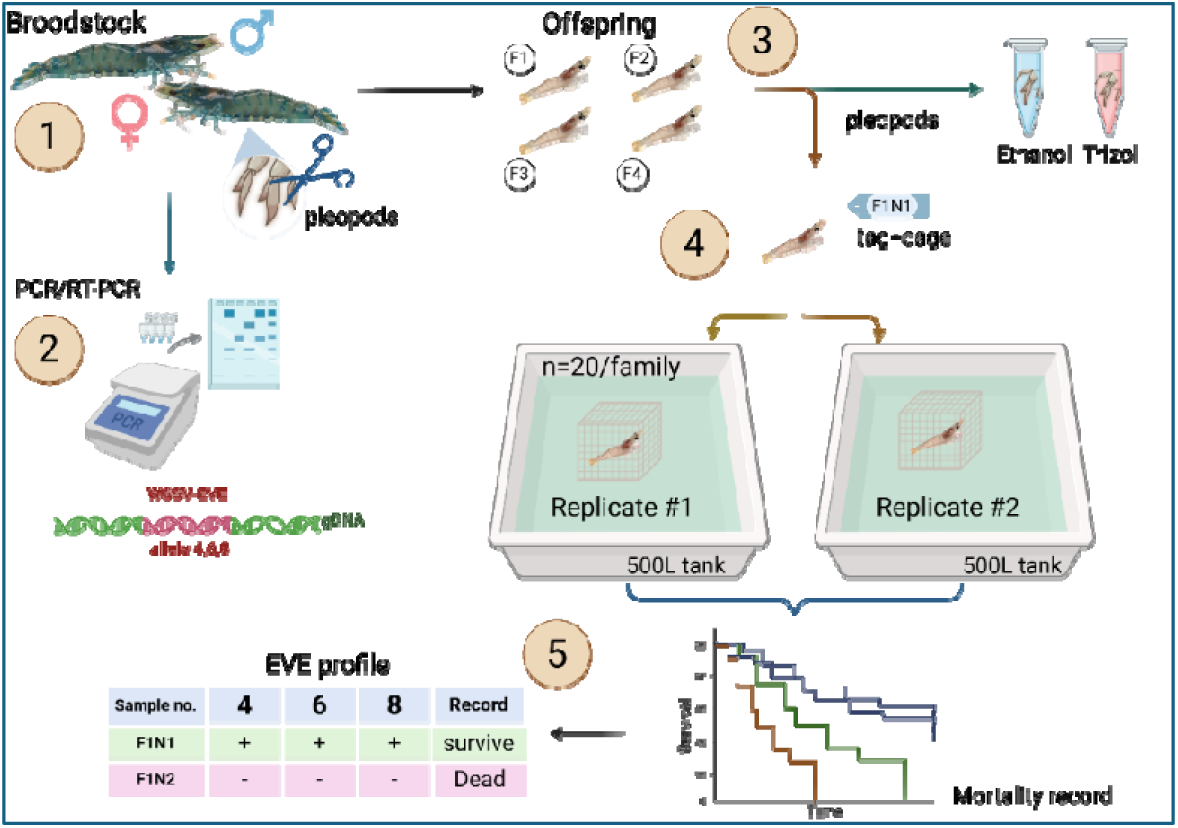
Schematic overview of the experimental design to evaluate the correlation between WSSV-EVE profiles and shrimp survival. (1) Pleopod tissue samples are collected from male and female *Penaeus monodon* broodstock. (2) Genomic DNA (gDNA) and RNA were extracted and screened using PCR/RT-PCR to determine the presence of specific WSSV-EVE alleles (e.g., alleles 4, 6, 8). (3) Offspring from various families (F1–F4) were sampled for pleopods and preserved in ethanol or Trizol for genotyping. (4) Individual shrimp from each family were tagged and caged, then distributed into two replicate 500 L tank systems (n=20 per family per tank). (5) EVE genotyping and its transcript profiles were matched with survival data.

### 2.3. Nucleic acid extraction

Genomic DNA and total RNA were extracted from pleopod tissues collected from both broodstock and selected offspring. Genomic DNA was isolated using the Exgene™ Cell SV DNA Extraction Kit (GeneAll, Korea) according to the manufacturer’s instructions. Total RNA was extracted using the chloroform–isopropanol extraction method. To eliminate genomic DNA contamination, RNA samples were treated with 20 U of DNase I (New England Biolabs, USA) at 37LJ°C for 1 hour, followed by re-precipitation using isopropanol. The concentrations of both DNA and RNA were determined using a NanoDrop spectrophotometer (Thermo Fisher Scientific, USA), and the samples were diluted to a final concentration of 50 ng/μL for downstream analyses.

### 2.4. PCR analysis of WSSV-EVE in broodstock and offspring

Three PCR and RT-PCR primer sets arbitrarily named methods 1 to 3 (**Table 1**) were designed based on the high frequency read sequences (HFRS) of EVE mapped to the WSSV genome (Tangchaiyaphum et al. 2024). Pleopod DNA from each broodstock specimen was used as the template and four female-male broodstock pairs were selected based on WSSV-EVE profiles for positive or negative EVE and for matching RNA expression (positive or negative). The same primers were used to detect EVE and their expressions with nucleic acids extracted from pleopods removed from the offspring prior to WSSV challenge. The table also includes the PCR primers used for virus quantification post challenge and for sex determination.

**Table 1.**
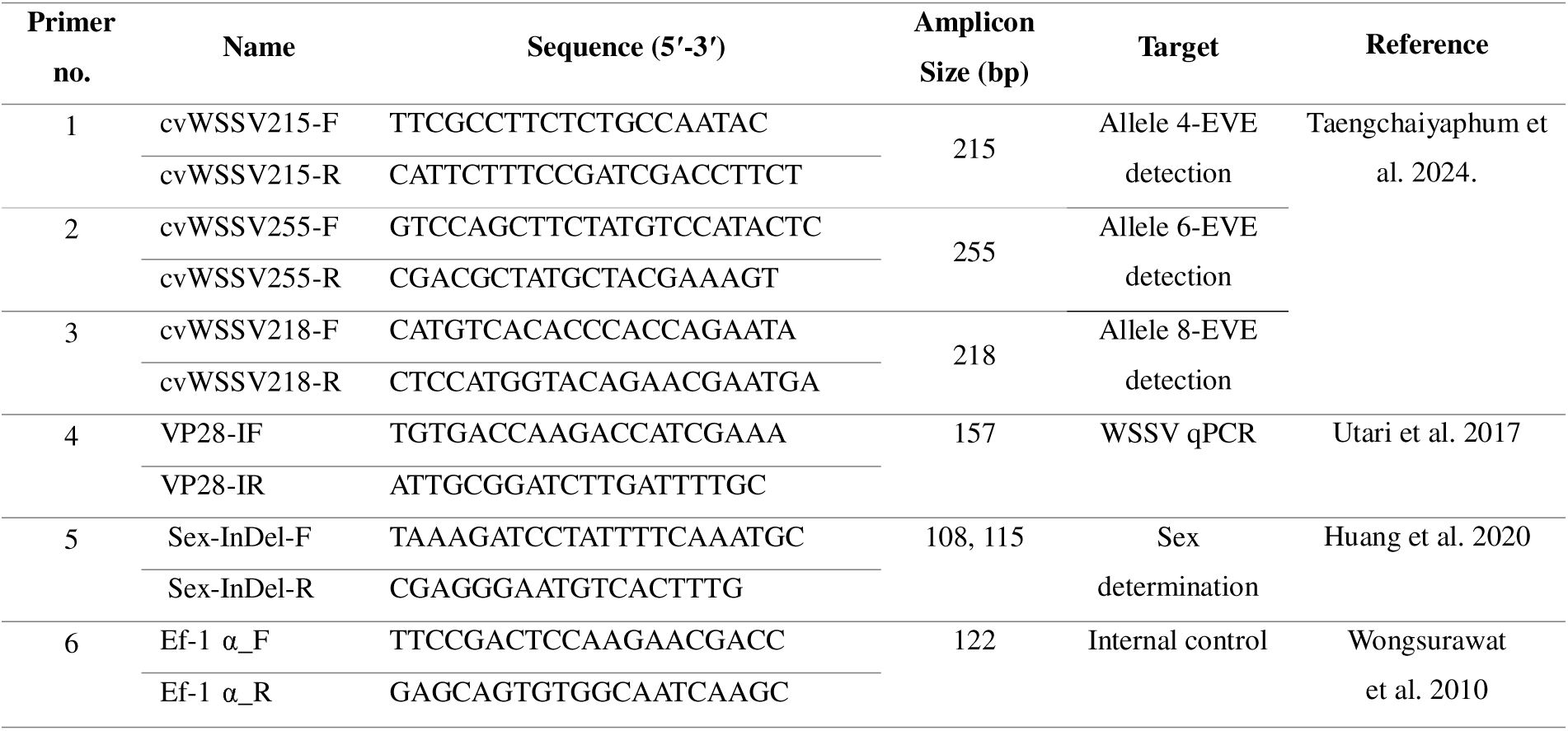
List of PCR and RT-PCR primers used in this study.

Based on the previously observed HFRS fragments in *P. vannamei* (Taengchaiyaphum et al., 2024), PCR analysis of HFRS fragments in *P. monodon* DNA samples taken prior to WSSV challenge were carried out to determine the correlation between WSSV-EVE profile and virus tolerance capacity in *P. monodon*. A total of 100 ng of genomic DNA was subjected to PCR amplification using the OneTaq^®^ PCR Master Mix (New England Biolabs, USA). The reaction mixture contained 1× PCR Master Mix, 0.4 μM each of forward and reverse primers, and 100 ng of template DNA. The thermal cycling protocol included an initial denaturation at 94 °C for 2 min, followed by 35 cycles of denaturation at 94 °C for 30 sec, annealing at 55 °C for 30 sec, and extension at 72 °C for 30 sec. A final extension step was performed at 72 °C for 5 min. PCR products were resolved by electrophoresis on a 1.5% agarose gel and visualized by ethidium bromide staining. Gel imaging and analysis were performed using the EZ Gel Documentation System (Bio-Rad, USA). Reverse transcription PCR (RT-PCR) analysis was performed to detect EVE transcripts corresponding to EVE-DNA signals observed in individual samples. The reaction was conducted using the SuperScript™ III One-Step RT-PCR System (Invitrogen, USA). Each reaction mixture contained 1× PCR Master Mix, 0.4 μM of each forward and reverse primer, and 100 ng of total RNA as template. The thermal cycling protocol involved reverse transcription at 50 °C for 20 min, followed by initial denaturation at 94 °C for 2 min, then 35 amplification cycles consisting of denaturation at 94 °C for 30 sec, annealing at 55 °C for 30 sec, and extension at 72 °C for 30 sec. A final extension step was performed at 72 °C for 5 min. Amplified products were resolved by electrophoresis on a 1.5% agarose gel and visualized by ethidium bromide staining.

PCR-based sex determination was conducted using the Sex InDel primer set as described by Huang et al. (2020). The PCR reactions were performed under the same conditions previously described, with the exception of a reduced annealing temperature set to 41 °C for 30 sec. Sex was determined based on the size of the amplified product: a 108 bp fragment indicated a male genotype, while a 115 bp fragment indicated a female genotype.

### 2.5. Selection of broodstock pairs to produce offspring for WSSV challenge

Altogether, 4 crosses were analyzed using females and males with HFRS profiles for putative EVE arbitrarily numbered 4, 6 and 8, with zero indicating absence of one of these alleles while a number indicates presence of a particular EVE 4, 6, 8 (**Table 2**). In addition, black typeface indicates that the particular EVE has no RNA expression while red typeface indicates RNA expression, although the sense of expression (i.e., positive, negative or dual) was not determined.

**Table 2.** Profiles of the 8 broodstock specimens selected for 4 crosses to produce offspring for use in challenge tests with WSSV. Red bold typeface indicates RNA expression while black indicates no expression and zero indicates absence of the relevant EVE.

| Cross # | Female no. | EVE profile | Male no. | EVE profile |
| --- | --- | --- | --- | --- |
| 1 | F13 | 0,0,0 | M47 | **4,6,8** |
| 2 | F98 | 0,0,0, | M106 | **4,6,8** |
| 3 | F38 | **4,6,8** | M17 | **4,0,0** |
| 4 | F64 | **4,6,8** | M74 | **4,6,8** |

### 2.6. WSSV challenge

A total of 40 juvenile shrimp (2-3 g) from each cross were divided into 2 replicates of 20 each for WSSV challenge test. They were individually tagged and 2 pleopods were removed from each, labeled and preserved in 95% ethanol for DNA and Trizol^®^ reagent (Invitrogen, USA) for RNA analysis. Each individual was housed in its own cage within the 500 L challenge tank.

The crude virus was prepared from an infected shrimp. In brief, WSSV-infected shrimp were prepared collected, and the whole body except the hepatopancreas was minced and homogenized in 1x PBS, pH 7.4. Tissue homogenate was filtered through a nylon net and the homogenate was aliquoted into 50 ml tubes. The virus stock was kept in -80°C and the concentration was determined by qPCR. For immersion challenge, the crude homogenate was poured into a 500 L challenge tank to obtain a final concentration at 10^6^ copies/ml seawater. Shrimp mortality was recorded daily for 14 days. Pleopods from each dead/moribund and surviving shrimp were collected for DNA extraction followed by qPCR to determine WSSV viral loads in copies per ng DNA.

### 2.7. Virus quantification in challenged offspring

WSSV loads were quantified by qPCR in DNA extracts to compare WSSV loads between dead and surviving shrimp for additional comparison with their EVE profiles. The primers used were VP28-IF and VP28-IR (Table 1). Ten nanograms of DNA was subjected to qPCR by using KAPA SYBR^®^ FAST qPCR Master Mix (Roche) and 40 cycles of PCR amplification was performed by CFX96™ Real-Time PCR Detection System (Bio-Rad, USA) following the protocol: 95 °C for 15 s followed by 40 cycles of 95 °C for 15 s, 55 °C for 30 s and 72 °C for 30 s. This was followed by a dissociation stage of 95 °C for 15 s, 60 °C for 1 min, 95 °C for 15 s and 60 °C for 1 min. All reactions were run with two replicates. The recombinant plasmid harboring virus fragments (VP28 gene) was applied as a standard to calculate copy numbers for each sample.

### 2.8. Preparation, sequencing and analysis of cvcDNAs from the list of dead and surviving offspring post virus challenge

To evaluate the correlation between cvcDNA profiles and WSSV tolerance, comparative detection of cvcDNA reads in offspring within the same family showing either low (dead) or high (survivor) tolerance to WSSV infection was evaluated using DNA isolated prior to WSSV challenge. The offspring of Cross 1 (F64 x M74) were selected and the DNA extracts of 10 individuals of dead and surviving lists were arbitrarily collected for circular DNA extraction. The method of circular DNA extraction from shrimp genomic DNA using plasmid-safe DNase digestion was as previously described (Taengchaiyaphum et al. 2024). Briefly, 2LJµg of DNA was incubated with 20LJU of plasmid-safe DNase (Epicentre, UK) at 37 °C for 4 days to digest linear DNA, with an additional 20LJU of enzyme added every 24 h. Remaining circular DNA was precipitated with absolute ethanol, pelleted by centrifugation, washed with 75% ethanol, and resuspended in PCR-grade water. Remaining circular DNA concentration was measured the using a Qubit Fluorometer with the broad range dsDNA assay kit (Invitrogen, USA). Subsequently, 40 ng of circular DNA (pooled from 10 shrimp of each dead or surviving list) was subjected to rolling circle amplification (RCA) using the REPLI-g Mini Kit (Qiagen, Germany). The RCA product was precipitated with ethanol, resuspended in PCR-grade water, and quantified using a NanoDrop spectrophotometer. A total of 20 µg of RCA product was submitted for DNA sequencing.

Sequencing of the rolling circle amplification (RCA) product was performed and analyzed by Novogene (Novogene Corporation Inc., Singapore) as previously described (Taengchaiyaphum et al. 2024). Briefly, 20 µg of RCA product was randomly fragmented by sonication. The resulting DNA fragments were end-polished, A-tailed, and ligated to full-length Illumina adapters, followed by PCR amplification using P5 and indexed P7 primers. The amplified libraries were purified using the AMPure XP system (Beckman Coulter, USA), assessed for size distribution using the Agilent 2100 Bioanalyzer (Agilent Technologies, CA, USA), and quantified by real-time PCR to ensure a minimum concentration of 3 nM. For data analysis, low-quality reads were filtered to obtain clean reads.

To identify viral sequences within pooled circular DNA reads, the viral DNA read analysis platforms followed a previously described method reported by Taengchaiyaphum et al. (2024). The raw reads were first subjected to quality control processing, including trimming and filtering using Trimmomatic, followed by quality assessment with FastQC. Putative viral sequences were identified using DIAMOND, a sensitive protein-based sequence alignment tool (Vilsker et al., 2019). Reads exhibiting ≥97% identity to a reference virulent strain of white spot syndrome virus (WSSV, GenBank accession no. AF440570) were mapped to the WSSV genome to determine their distribution across the viral genome.

### 2.9. Mapping of putative WSSV-EVE reads from pooled *P. monodon* cvcDNA

To identify viral sequences within pooled circular DNA reads, two viral DNA read detection platforms were employed. The first approach followed a previously described method reported by Taengchaiyaphum et al. (2024), which was based on proprietary analysis by a private company. The second method utilized the Genome Detective platform (available online at http://www.genome-detective.com/app/typingtool/virus/), as described in a recent publication. For both methods, paired-end reads in FASTQ format were used. In the Genome Detective platform, raw reads were first subjected to quality control processing, including trimming and filtering using Trimmomatic, followed by quality assessment with FastQC. Putative viral sequences were identified using DIAMOND, a sensitive protein-based sequence alignment tool (Vilsker et al., 2019). Reads exhibiting ≥97% identity to a reference virulent strain of white spot syndrome virus (WSSV, GenBank accession no. AF440570) were mapped to the WSSV genome to determine their distribution across the viral genome.

## 3. RESULTS

### 3.1. Overall survival of offspring in the 4 WSSV challenges

The offspring from 4 crosses were subjected to WSSV challenge for 14 days. The survival results are shown in **Figure 2**. All offspring from Cross 1 (F64 x M74) and Cross 2 (F38 x M17) died, while mean survival in Cross 3 (F13 x M47) and Cross 4 (F98 x M106) were 60.0±20.0% and 47.2±8.3%, respectively and was associated with presence and expression of EVE-4,6,8. Details of the analysis for each cross are given in the following sections.

**Figure 2:**
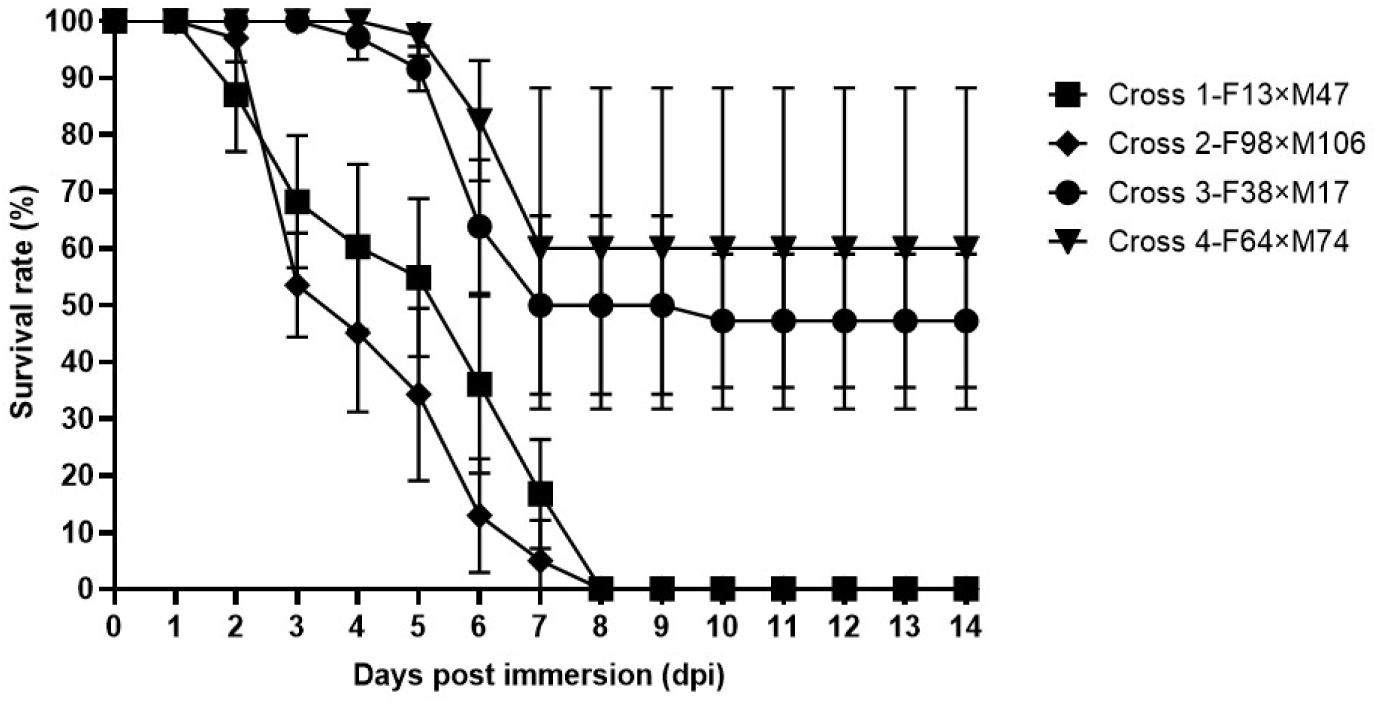
The survival rates of the offsprings from 4 crosses post WSSV challenge. The number of 40 offsprings from each cross were subjected to WSSV challenge by immersion method. The survival of the offsprings were recorded for 14 days.

**Figure 2.**
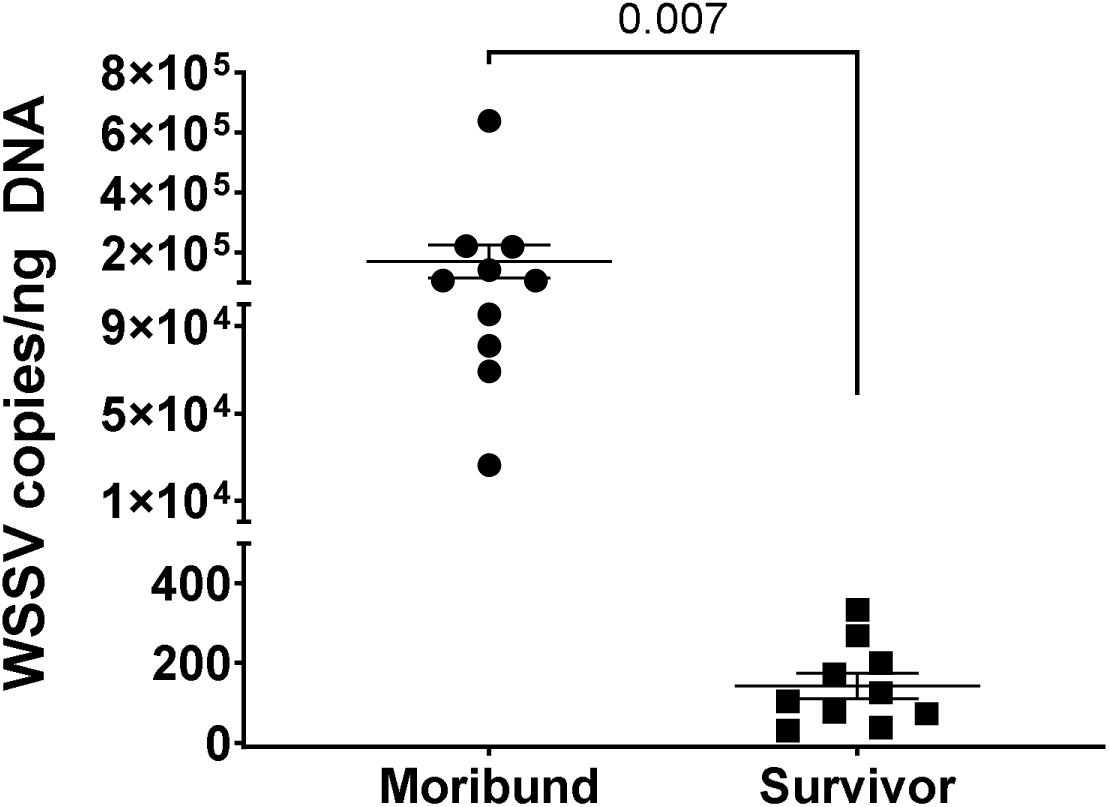
Quantitative PCR (qPCR) analysis showing very high WSSV loads in dead shrimp and very low WSSV loads in surviving shrimp from the WSSV challenge test. Shrimp specimens were from the offsprings of Cross 1 (F64XM74).

### 3.2. Detailed outcomes of WSSV challenges using offspring from 4 crosses

#### 3.2.1. Results from WSSV challenge with offspring from Cross 1: F13 x M47

The overall results from the challenge of 40 juvenile shrimp offspring from a cross between F13 and M47 are shown in **Table 3**. The absence of an EVE is indicated by a black zero (0) while presence without expression is indicated by the EVE number in bold black (e.g., **4**). In contrast, possession of an EVE together with its RNA expression is indicated as a bold, red number (e.g., **4**). However, the positive, negative or dual sense of RNA expression was not determined. Sex was determined by PCR since the shrimp were too small for sexing. There were two challenge groups comprising 20 shrimp, each in separate aquaria. All shrimp died within 9 days after challenge. To analyze for the presence of EVE, 10 specimens were arbitrarily selected. These were 6 from Replicate #1 and 4 from Replicate #2. Note that sex could not be determined physically but was done by PCR, so exact selection of 5 males and 5 females for analysis was not possible.

**Table 3.** Results from WSSV challenge of offspring from Cross 1: F13 x M47. *Note: One shrimp died following pleopod removal in Replicate #2.

| Day post | Number of dead shrimp |
| --- | --- |

| Immersion | Replicate # 1<br>(samples no. 1-20) | Replicate # 2<br>(samples no. 21-40) |
| --- | --- | --- |
| 1 | 0 | 0 |
| 2 | 4 | 1 |
| 3 | 4 | 3 |
| 4 | 2 | 1 |
| 5 | 1 | 1 |
| 6 | 4 | 3 |
| 7 | 3 | 4 |
| 8 | 2 | 4 |
| 9 | 0 | 2 |
| **Total** | **20** | **19\*** |

The results of EVE analysis for Cross 1 are shown in **Table 4**. In the 10 arbitrarily selected shrimp specimens, there were 7 females and 3 males instead of 5 each. However, with the small sample size, 7:3 and 5:5 were not significantly different by the Fisher Exact test (p = 0.32). Parental Female F13 was PCR negative for EVE-4,6,8 while Male M47 was positive for all 3 alleles together with RNA expression for all. The positive, negative or dual sense of these expressions were not determined. In the table, absence of an EVE is indicated by a bold, black number **0** while presence without expression is indicated by the EVE number in bold black (e.g., **4**). In contrast, possession of an EVE together with its RNA expression is indicated as a bold, red number (e.g., **4**). Because all the shrimp died, only 10 of the tagged shrimp were arbitrarily selected for detailed PCR analysis of their DNA from the pleopod samples taken before the challenge. Sex was also determined by PCR since the shrimp were too small for sexing. Thus, the proportions of males and females in all the crosses described herein could not be uniformly collected as half male and half female.

**Table 4.** Results of Cross 1 (F13 x M47) EVE analysis of DNA from 10 arbitrarily selected offspring, all of which died in the WSSV challenge test. Red typeface indicates RNA expression. Gray background indicates males and white females. Note: **f**; female, **m**; male.

| Parents | Allele 4 | Allele 6 | Allele 8 |
| --- | --- | --- | --- |
| F13 | 0 | 0 | 0 |
| M47 | 4 | 6 | 8 |
| Offspring no. |  |  |  |
| 4f | 0 | 6 | 8 |
| 14f | 0 | 6 | 8 |
| 18f | 0 | 6 | 0 |
| 28f | 0 | 0 | 0 |
| 3f | 4 | 6 | 0 |
| 12f | 0 | 6 | 0 |
| 29f | 0 | 6 | 0 |
| 19m | 4 | 6 | 0 |
| 23m | 0 | 6 | 0 |
| 40m | 0 | 6 | 0 |
| Transmitted | 2 | 9 | 2 |
| Expressed | 1 | 2 | 0 |

It can be seen in Table 4 that none of the 10 offspring carried all three EVE-4 -6 and -8, so it is unlikely that these alleles were linked together on a single chromosome in M47. If all 3 had been linked, it would have been predicted that 50% of the 10 offspring (5 shrimp) would have inherited all three EVE. However, we see no shrimp that inherited all three.

Only 1 shrimp specimen (28f) was not positive for any of the 3 EVE from M47. The Mendelian probability for such an outcome would be 0.5 x 0.5 x 0.5 = 0.125, which multiplied by 10 would predict 1 specimen in 10 randomly selected individuals. For the small number of 10 shrimp tested for a single allele, only a ratio of 0:10 would have been significantly different from a ratio of 5:5 by the Fisher exact test. Thus, we concluded that each of **EVE-4, -6** and **-8** was transmitted independently to the offspring at a ratio of 1:1 (50%), that would amount to 5 predicted offspring out of 10 for each EVE. As it turned out, the ratios were 2:8 in10 specimens for both **EVE-4** and **EVE-8,** and 9:1 specimens for **EVE-6**.

Examining the issue of 9 in 10 specimens giving positive test results for transmission of **EVE-6**, we considered the possibility that **EVE-6** in M47 might have been present in two copies, but with each heterozygous and each on a different chromosome. As shown in the Punnett square in **Table 5**, this would have given 4 possible gamete combinations for the dual-positive parent (**6a** + **6b**; **6a** + 0b; **6b** + 0a; and 0a + 0b). When crossed with a negative parent, the result would be a ratio of 3:1 (75%) for offspring receiving allele 6 in a single or double state and with 2/3 of the **EVE-6**-positive offspring showing expression and 1/3 not. However, the two of these possibilities could not be distinguished by statistical analysis due to the low number of samples.

**Table 5.** Punnett square of the possible distribution if two copies of **EVE-6** (here illustrated as **EVE-6a** and **EVE-6b**) were present in M47, each heterozygous and on different chromosomes. The unexpressed **EVE-6b** would not have been discoverable in the analysis of M47.

| **F13** | **M47** |  |  |  |
| --- | --- | --- | --- | --- |
|  | **6a + 6b** | **6a + 0b** | **6b + 0a** | **0a + 0b** |
| **0a + 0b** | 6a + 6b + 0a + 0b | 6a + 0b + 0a + 0b | 6b + 0a + 0a + 0b | 0a + 0b + 0a + 0b |
| **0a + 0b** | 6a + 6b + 0a + 0b | 6a + 0b + 0a + 0b | 6b + 0a + 0a + 0b | 0a + 0b + 0a + 0b |

An important feature of Cross 1 was the expression of all 3 **EVE-4,6,8** in parental M47 but expression in a single EVE in one of two offspring specimens carrying **EVE-4** and two of nine specimens carrying **EVE-6** and no expression in two specimens carrying **EVE-8**. Clearly, there were additional controls over EVE expression that are inherited from mated parents independently of the EVE.

#### 3.2.2. Outcome of WSSV challenge with offspring from Cross 2: F98 x M106

The results of the WSSV challenge test using offspring from Cross 2 are shown in **Table 6**. All the shrimp died within 8 days after challenge.

**Table 6.** Results from WSSV challenge of offspring from Cross 2: F98 x M106. *Note: 3 shrimp died following pleopod removal.

| **Day post**<br>**Immersion** | **Number of dead shrimp** |  |
| --- | --- | --- |
|  | **Replicate # 1**<br>**(samples no. 1-20)** | **Replicate # 2**<br>**(samples no. 21-40)** |
| 1 | 0 | 0 |
| 2 | 1 | 0 |
| 3 | 8 | 8 |
| 4 | 2 | 1 |
| 5 | 2 | 2 |
| 6 | 3 | 5 |
| 7 | 1 | 2 |
| 8 | 0 | 2 |
| **Total** | **17\*** | **20** |

The results of EVE analysis for 10 shrimp arbitrarily selected, (5 from Replicate #1 and 5 from Replicate #2) from Cross 2 are shown in **Table 7**. The female parent F98 was PCR negative for EVE-4,6,8 while the male M106 was positive for all 3 alleles but there was RNA expression in only 1 shrimp specimen each for **EVE-4** and **EVE-8**, but the plus or minus sense of RNA expression was not determined.

**Table 7.** Results for 10 offspring arbitrarily selected for EVE analysis from Cross 2 (F98 x M106). All the challenged offspring died. Alleles shown in red typeface were expressed while those in black were unexpressed. Gray background indicates males and white females. **f**; female, **m**; male

| Parents | Allele 4 | Allele 6 | Allele 8 |
| --- | --- | --- | --- |
| F98 | 0 | 0 | 0 |
| M106 | 4 | 6 | 8 |
| Offspring |  |  |  |
| 1f | 0 | 6 | 0 |
| 2f | 0 | 6 | 0 |
| 4f | 4 | 6 | 0 |
| 29f | 0 | 0 | 0 |
| 30f | 0 | 0 | 0 |
| 31f | 0 | 6 | 0 |
| 21f | 0 | 6 | 0 |
| 5m | 0 | 6 | 0 |
| 7m | 0 | 6 | 8 |
| 27m | 0 | 0 | 0 |
| Transmitted | 1 | 7 | 1 |
| Expressed | 1 | 1 | 0 |

In summary for Cross 2, the overall outcome was similar to that for Cross 1. The three EVE were inherited independently in Mendelian fashion, but RNA expression did not always correspond to that of the parent from which it originated. Thus, the results, like those in Cross 1, indicated that control of EVE expression was inherited independently of the EVE, and that the ratio of 1:10 did not differ significantly from 5:5 due to the low sample number. In addition, the argument for a second, independent, EVE-6 allele in the parent M106 was also possible (e.g., Table 4 for Cross 1).

Also similar to Cross 1, there was no indication of linkage between alleles **EVE-4, 6** and **8**. In addition, the distribution of 7 females to 3 males instead of 5 to 5 was not significantly different due to the small number of 10 specimens. The reason for total mortality may be that EVE in only 2 specimens showed RNA expression, and/or that expression might have been in positive instead of negative sense. On the other hand, it is also possible that there might be additional genetic control(s) regarding the protective ability of an antisense RNA transcript. As in Cross 1, the results indicated that control over EVE expression and function may be governed by one or more unknown, additional and currently unknown factors.

#### 3.2.3. Outcome of WSSV challenge with offspring from Cross 3: F38 x M17

The results of the WSSV challenge test using offspring from Cross 3 are shown in **Table 8**. From two replicates of 20 shrimp each, there were 11 dead shrimp in Replicate 1 and 8 in replicate 2, resulting in a mean survival of 11 in 20 (55%).

**Table 8.** The results of the WSSV challenge test using offspring from Cross 3: F38 x M17.

| Day post<br>Immersion | Number of dead shrimp |  |
| --- | --- | --- |
|  | Replicate # 1<br>(samples no. 1-20) | Replicate # 2<br>(samples no. 21-40) |
| 1 | 0 | 0 |
| 2 | 0 | 0 |
| 3 | 0 | 0 |
| 4 | 0 | 1 |
| 5 | 1 | 1 |
| 6 | 7 | 3 |
| 7 | 3 | 2 |
| 8 | 0 | 0 |
| 9 | 0 | 0 |
| 10 | 0 | 1 |
| **Total** | **11** | **8** |

The results of EVE analysis for Cross 3 are shown in **Table 9**. PCR revealed that there were 9 females and 11 males in the arbitrarily collected 10 dead shrimp (7 from Replicate #1 and 3 from Replicate #2) and 10 surviving shrimp (7 from Replicate #1 and 3 from Replicate #2). This was not significantly different from the expected 10 to 10 by Fisher Exact test (p = 0.5). The parental female F38 was PCR positive for the presence of **EVE-4,6,8** and all were expressed, while the male M17 was positive for expressed **EVE-4** only. As in Crosses 1 and 2, the sense of EVE expression was not determined, but unlike Crosses 1 and 2, above, 20 shrimp were analyzed instead of 10. This was because there were both survivors and dead shrimp after the WSSV challenge and it was of interest to determine whether there was any relationship between survival and presence of expressed EVE.

**Table 9.**
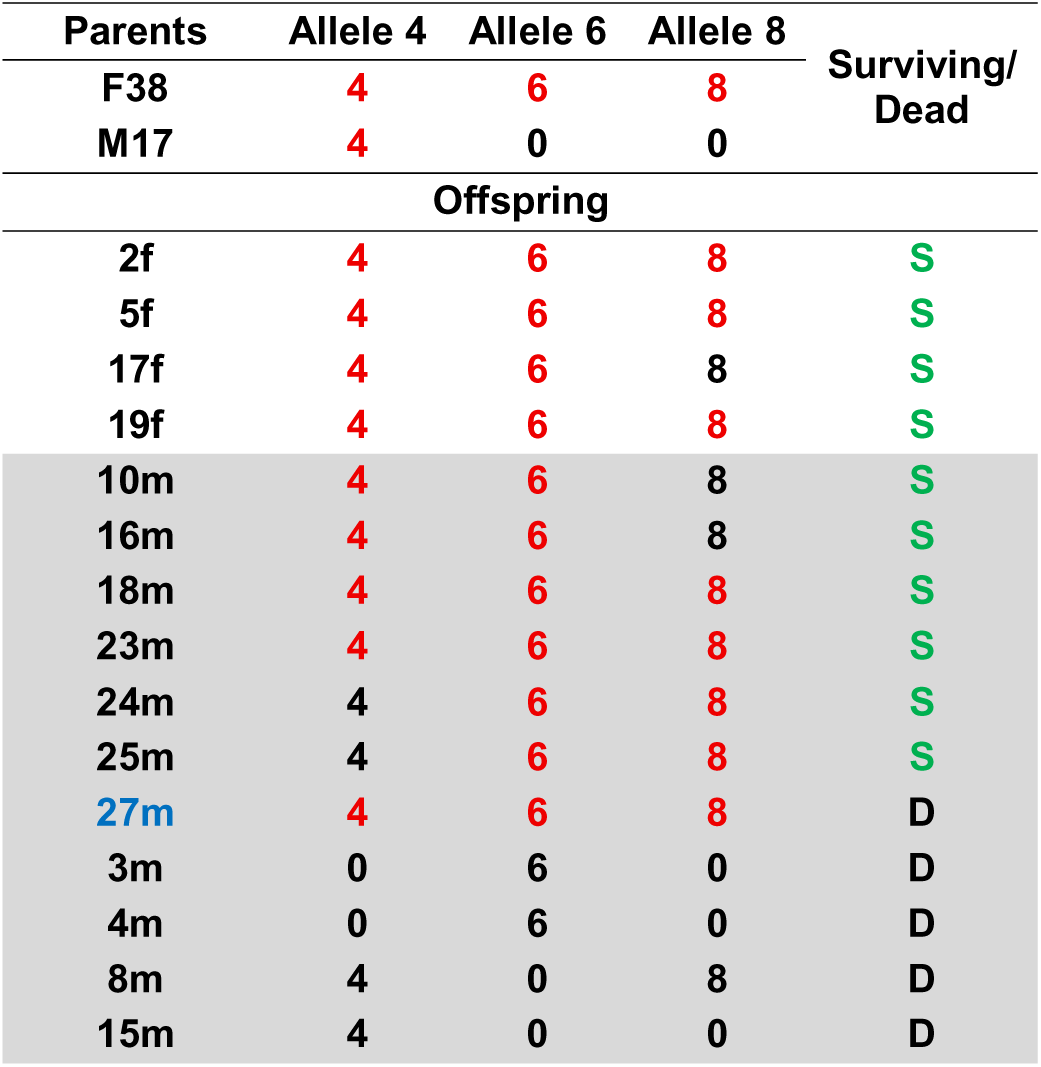
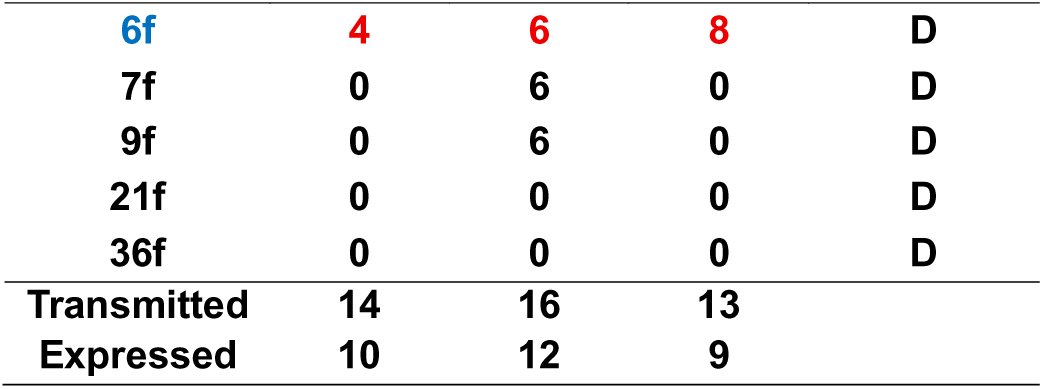
Analysis of the offspring from Cross 3 (F38 x M17). Gray shading highlights males and white females. Surviving (S) and dead (D) shrimp are indicated in green and black typeface, respectively. The offspring numbers in blue typeface represent samples with unexpected results because they died despite RNA expression of all three EVE-4,6,8.

| Parents | Allele 4 | Allele 6 | Allele 8 | Surviving/<br>Dead |
| --- | --- | --- | --- | --- |
| F38 | 4 | 6 | 8 |  |
| M17 | 4 | 0 | 0 |  |
| Offspring |  |  |  |  |
| 2f | 4 | 6 | 8 | S |
| 5f | 4 | 6 | 8 | S |
| 17f | 4 | 6 | 8 | S |
| 19f | 4 | 6 | 8 | S |
| 10m | 4 | 6 | 8 | S |
| 16m | 4 | 6 | 8 | S |
| 18m | 4 | 6 | 8 | S |
| 23m | 4 | 6 | 8 | S |
| 24m | 4 | 6 | 8 | S |
| 25m | 4 | 6 | 8 | S |
| 27m | 4 | 6 | 8 | D |
| 3m | 0 | 6 | 0 | D |
| 4m | 0 | 6 | 0 | D |
| 8m | 4 | 0 | 8 | D |
| 15m | 4 | 0 | 0 | D |
| 6f | 4 | 6 | 8 | D |
| 7f | 0 | 6 | 0 | D |
| 9f | 0 | 6 | 0 | D |
| 21f | 0 | 0 | 0 | D |
| 36f | 0 | 0 | 0 | D |
| Transmitted | 14 | 16 | 13 |  |
| Expressed | 10 | 12 | 9 |  |

Of the 20-shrimp selected for analysis, 9 were identified as females and 11 as males which is not significantly different (p = 0.5) from the expected Mendelian distribution of 10:10 by the Fisher Exact test. We can conclude that female F38 positive for “alleles” 4, 6 and 8 could not have been homozygous for one or more of the three alleles. Otherwise, all the offspring would have been positive for whichever independent allele was homozygous. Thus, we can conclude that the female was heterozygous for all three alleles and that the probability of offspring receiving each allele from this parent would be 0.5 or 50% by Mendelian distribution. However, the male M17 also carried allele 4, so the probability of getting that allele from both the male and female would increase the total probability of getting at least 1 copy of allele to 75%, yielding a predicted distribution of 15 specimens positive and 5 negative for allele 4. The actual numbers obtained were 14 positive and 6 negatives, which was not significantly different from the expected distribution by Fisher Exact test (p = 0.5).

For allele 6, the predicted distribution was 10:10 (50%) but the actual distribution was 16 positive (80%) and 4 negative, and this was significantly different (p = 0.05) by the Fisher Exact test. This supported the proposal that an additional allele 6 was present as an independent allele at another location in the shrimp genome (see Table 3) that would yield an expected distribution of 3:1 or 15 to 5 which was not significantly different (p = 0.5) from actual 16:4 by the Fisher Exact test. In addition, the result for EVE-8 was 13:7 (65%) but was not significantly different from the expected 10:10 by the Fisher Exact test (p = 0.26). These results suggest that larger samples are needed to clearly determine the possibility of 2 or more independent EVE of the same sequence.

With respect to the predicted probability for an individual offspring to receive all 3 of the unlinked alleles in this cross would have been 0.75 x 0.5 x 0.5 = 0.188 which would be about 3-4 specimens. Instead, we found the number of offspring carrying all 3 alleles was 12/20 = 60%. The simplest explanation was that the 3 “alleles” tended to be inherited together. This, in turn, suggested that they were linked on the same chromosome.

**Table 9** shows that out of 10 arbitrarily selected dead and living shrimp each, the distribution was 9 females was 11 males. In addition, the ratio of surviving females to males was 4:6, while that for dead shrimp was 5:5 indicating no bias for sex in the outcome of the test. However, in the group of 10 survivors, all carried expressed EVE of at least 2 out of the 3 alleles 4,5,6 while only 2 individuals that carried these three alleles were dead. Thus, out of 12 shrimp with this EVE profile the survival was 10/12 = 83%. In contrast, survival of the 8 shrimp that carried only 1 or none of these alleles was zero. An anomaly was that all 3 alleles were also expressed in 2 dead shrimp, (1 male and 1 female).

If the 3 alleles in the female parent were linked and present only one chromosome, then one would expect the number of offspring carrying all 3 alleles compared to the number with no alleles would be approximately 1:1 (10:10 in 20 samples). What was found was 12 in 20 with all three alleles and only 2 with no alleles. For the remaining 6 specimens, five showed 1 allele and one showed 2 alleles. The ratio of 12:8 was reasonably close to the expected Mendelian ratio of 10:10 and was not significantly different from it by the Fisher exact test (p = 0.38). However, the low number of 2 specimens with none of the three alleles may be lower than expected (i.e., the paired chromosome with none of the 3 EVE). On the other hand, 8 specimens showing expression of 0, 1 or 2 alleles suggested they may be potential crossover partners from the matching chromosome proposed to carry the linked EVE-468.

Another interesting feature of the results is that expression of the transmitted alleles did not always match that of F38 and M17 that both showed expression for all the alleles they carried. For example, for 14/20 samples that received allele 4, 10 were expressed while 4 were not. For allele 6, 17/20 were transmitted and 12 were expressed while 5 were not. For allele 8, 13 were transmitted but only 9 expressed. Furthermore, of the 7 offspring that received and expressed all three alleles, 5 specimens were survivors and 2 were dead shrimp (6f and 27m). Is it possible that the dead specimens did not express antisense-RNA from any of the 3 expressed EVE? In summary, transmission of the EVE was Mendelian but expression was variable, suggesting an unknown, independent genetic factor(s) controlling expression or not (i.e., production of RNA or not) and also RNA expression type (positive sense, negative sense or dual sense).

All the surviving shrimp from the WSSV challenge carried combinations of 2 or 3 expressed alleles of 4,6 or 8, while most of the dead shrimp (8/10) either lacked these alleles or carried them without expression. However, there were also 2 dead shrimp (6f and 27m) that carried all 3 expressed alleles. We propose that these 2 dead specimens may have expressed RNA of positive sense only (non-protective), while the other 8 expressed RNA of negative or dual sense (protective). Additional work is required to test these proposals.

A Punnett square (**Table 10**) illustrating transmission of a heterozygous allele 6 from one parent mated with a parent not carrying that allele, the distribution of allele 6 in the offspring would be 1:1 (50%). However, we see for allele 4, that having the male also heterozygous for allele 4 raises the probability of any offspring receiving allele 4 in 1 or 2 copies to be 75%. Thus, we cannot exclude the possibility that F38 may carry independent copies of these alleles on separate chromosomes that are inherited independently of alleles in the triplet linkage group 4-6-8. Of course, there is also probable presence of EVE other than 4,6,8 in the genome that were not detected or studied.

**Table 10.**
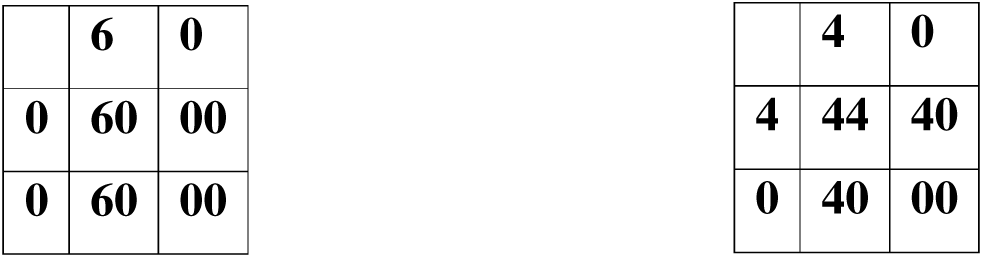
Punnett squares showing probabilities of occurrence of 50% offspring inheriting allele 6 present on one chromosome from 1 parent (50%) and 75% for inheriting at least 1 copy of allele 4 if it is present on one chromosome in each parent.

In Table 9, the distribution was 6 females to 14 males as determined by PCR, and this was not significantly different (p = 0.17) from the expected distribution of 10 to 10 using the Fisher Exact test. Of the arbitrarily selected 10 dead (9 from Replicate #1 and 1 from Replicate #2), there were 2 dead females and 8 dead males. Of the arbitrarily selected 10 survivors (4 from Replicate #1 and 6 from Replicate #2), there were 4 surviving females and 6 surviving males. The results indicated no bias for sex in the outcome of the test. However, in the group of 10 survivors, 9 carried expressed EVE of at least 2 out of the 3 alleles, while 4 other individuals carrying the same 2 alleles (Dead no. 18) or all 3 alleles (Dead no. 12, 19, 26) were dead. Thus, out of 13 shrimp with such EVE profiles the survival was 9/13 = 69%. In contrast, for the 6 shrimp that carried only 1 or none of these alleles, survival was zero. The anomalies were 3 dead shrimp that expressed all 3 alleles..

Altogether, this cross is of little use because of the difficulty in interpreting the results with common EVE present in both the male and female parents. A Punnett square for distribution of linked alleles 4,6,8 in the offspring of a cross with both parent heterozygous 4,6,8 and showing the prevalence would be 75%. However, the calculations are difficult because both parents carry all three alleles and it is possible that there may be other single copies, particularly for EVE-6, heterozygous on another chromosome and affecting the phenotypes of the offspring. It is also possible, if not likely, that there were other EVE in the shrimp that we do not know about but also affected survival. In the future, we should probably use crosses with all the EVE present in only one of the parents.

Overall in Cross 3, there was a clear association between survival and the presence of expressed EVE-4,6,8. Specifically, all 10 = 100% of the surviving shrimp carried and expressed two or three of these EVE. In contrast, 8/10 = 80% the moribund/dead shrimp showed either no EVE (2 shrimp) or 1 EVE without expression (5 shrimp) or 2 EVE without expression. The exception was 2 dead/moribund shrimp that expressed all 3 EVE. We propose that these latter 2 shrimp may indicate the existence of an unknown control or controls over the sense of RNA expression by EVE.

#### 3.2.4. Outcome of WSSV challenge with offspring from Cross 4: F64 x M74

The results of the WSSV challenge test using offspring from Cross 4 are shown in **Table 11**. The parental shrimp in this cross (F64 and M74) were both PCR-positive for EVE 4, 6 and 8 and all were positive for RNA expression of all 3 alleles. In two replicates of 20 shrimp each, there were 12 dead shrimp in Replicate 1 and 4 in replicate 2, for a mean survival of 12/20 (60%).

**Table 11.**
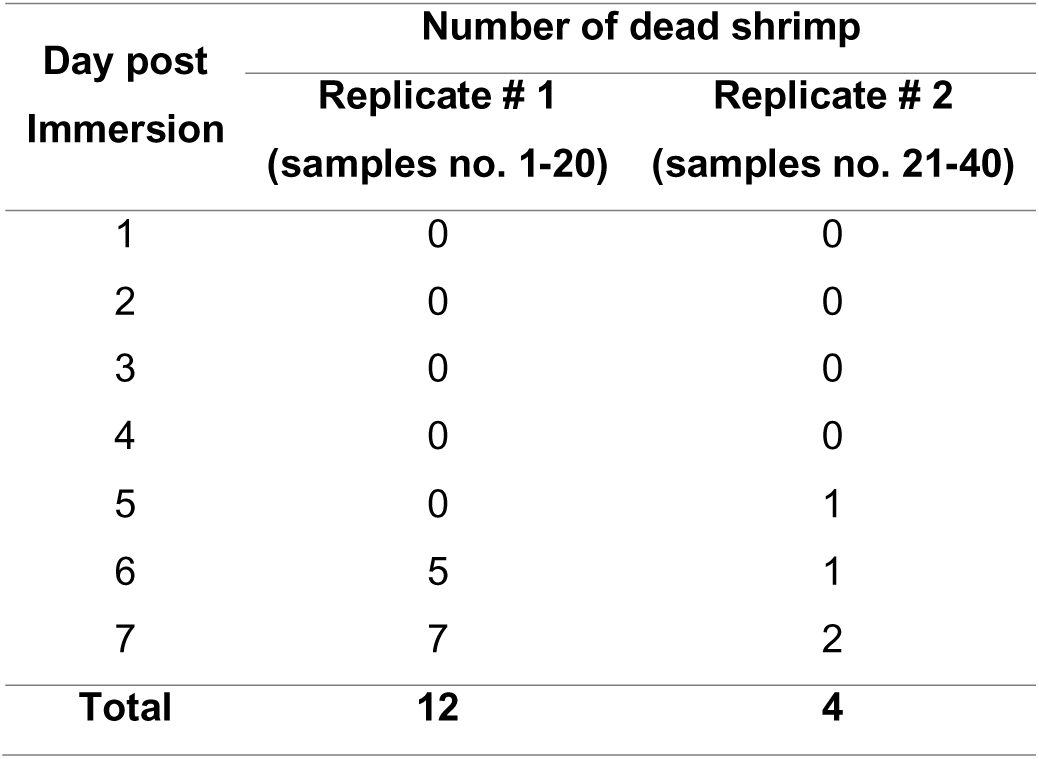
The results of the WSSV challenge test using offspring from Cross 4: F64 x M74.

| Day post<br>Immersion | Number of dead shrimp |  |
| --- | --- | --- |
|  | Replicate # 1<br>(samples no. 1-20) | Replicate # 2<br>(samples no. 21-40) |
| 1 | 0 | 0 |
| 2 | 0 | 0 |
| 3 | 0 | 0 |
| 4 | 0 | 0 |
| 5 | 0 | 1 |
| 6 | 5 | 1 |
| 7 | 7 | 2 |
| **Total** | **12** | **4** |

Results of testing for sex and EVE in offspring of Cross 4 using PCR with DNA extracted prior to WSSV challenge are shown in **Table 12**. When tested for sex the arbitrarily selected shrimp specimens consisted of 6 females and 14 males, while the expected ratio was 10 of each. The Fisher Exact test revealed that the observed ratio 6:14 was not significantly different from the expected ratio or 10:10 (p = 0.17). By PCR testing for EVE 4, 6 and 8, we could conclude that neither of the parents were homozygous for one or more of the three independent alleles, since no allele was positive in all 20 of the offspring. The frequency of each allele in the offspring was about the same 75-85% which coincided with the expected 75% from a cross of two shrimp each heterozygous for the same allele. The probability of getting all three alleles together in one individual would then be 0.75 x 0.75 x 0.75 = 42% which would be 8 specimens in 20. However, in our data we see 14/20 = 70% positive for all 3 alleles. As with Cross 3, the simplest explanation for this is that the 3 “alleles” tended to be inherited together, in turn, suggesting that they were linked together on the same chromosome, as in Cross 3. However, to allow for the variation in distribution to the offspring, they must be separated by sufficient distances to allow relatively high variation in the number transmitted to individual offspring due to crossover events during meiosis.

**Table 12.**
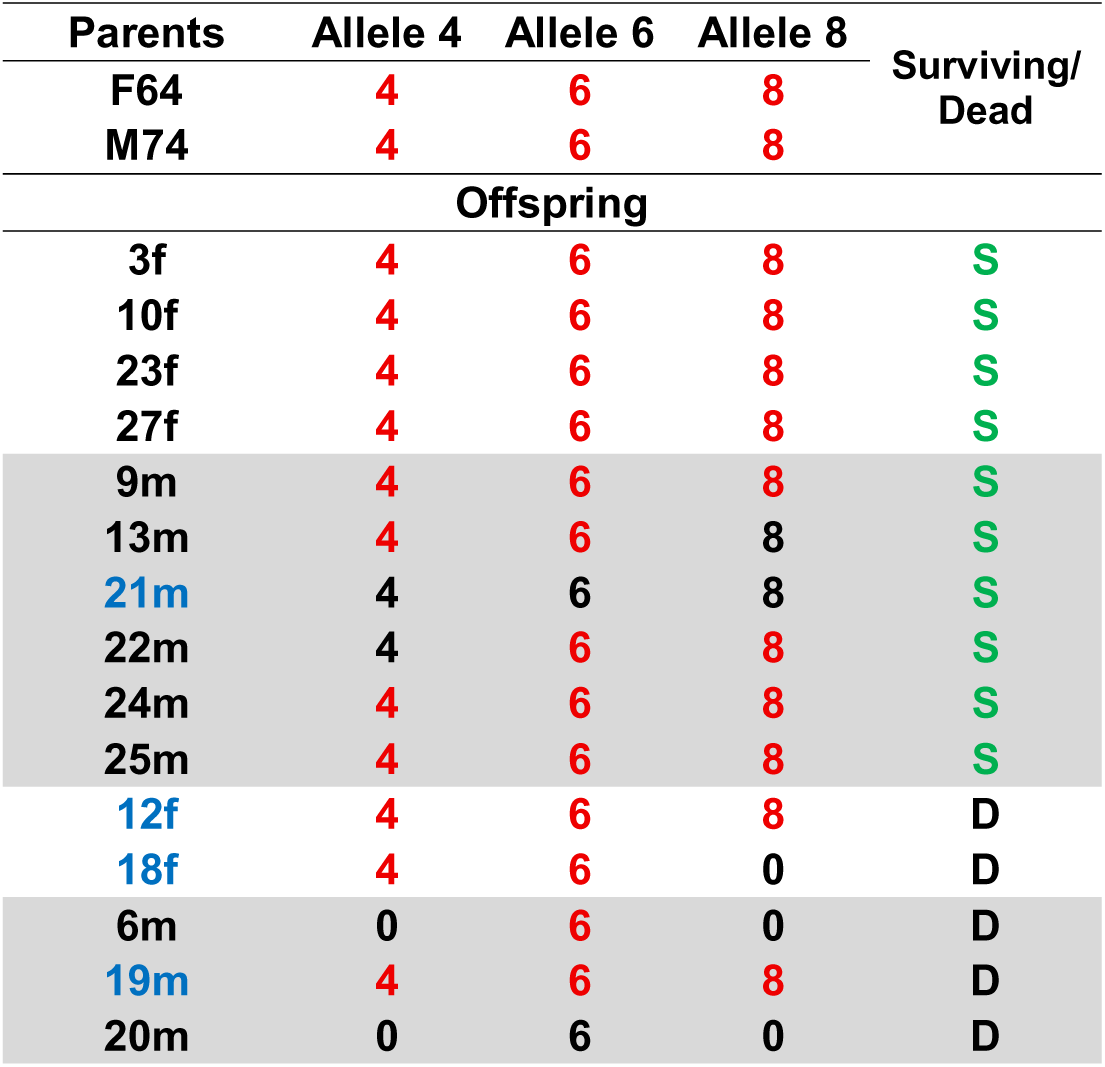
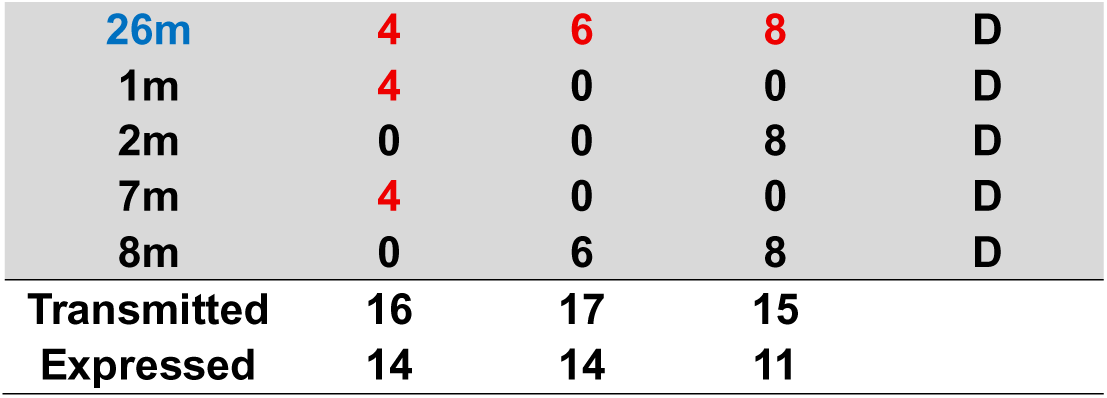
Analysis of the offspring from Cross 4 (F64 x M74) showing alleles transmitted with expression in red and without expression in black. Surviving (**S**) and Dead (**D**) shrimp are indicated in green and black text, respectively. The 4 specimen numbers in blue typeface indicate shrimp with unexpected phenotypes with respect to their genotypes.

In **Table 12**, the distribution was 6 females to 14 males as determined by PCR, and this was not significantly different (p = 0.17) from the expected distribution of 10 to 10 using the Fisher Exact test. In the group of arbitrarily selected 10 dead shrimp (9 from Replicate #1 and 1 from Replicate #2), there were 4 females and 6 males, and in the group of arbitrarily selected 10 surviving shrimp (4 from Replicate #1 and 6 from Replicate #2) there were (5 females and 5 males). This indicated no bias for sex in the outcome of the test. However, in the group of 10 survivors, 9 (90%) expressed either 2 or 3 alleles of EVE-4,6,8. In contrast, the 1 remaining survivor unexpectedly carried the same 3 alleles, but without expression, suggesting that one or more other protective EVE might be present. In contrast, mortality for the 6 shrimp that carried only 1 or none of these alleles, survival was zero. However, there were 3 dead shrimp that expressed all 3 of the EVE-4.6.8 alleles and 1 that expressed EVE-4,6. As in Cross 3, we surmise that these EVE did not express negative-sense RNA.

Altogether, this cross is of little use because of the difficulty to interpret results with common EVE present in both the male and female parents. A Punnett square for distribution of linked alleles 4,6,8 in the offspring of a cross with both parent heterozygous for linked 4,6,8 would be 75%. However, the calculations are difficult because both parents carry all three alleles and it is possible that there may be other single copies, particularly for EVE-6, heterozygous on another chromosome and affecting the phenotypes of the offspring. It is also possible, if not likely, that there were other EVE in the shrimp that we do not know about but also affect survival. In the future, it would be best to use crosses with all the EVE present in only one of the parents and the other negative for the same EVE.

### 3.3. qPCR analysis for WSSV loads in dead and surviving shrimp from Cross 4

When qPCR analysis of WSSV replication was performed on DNA extracted from arbitrarily selected 10 dead shrimp and 10 surviving shrimp from Cross 1 (F64xM74) above, the dead shrimp exhibited significantly higher viral loads (1.7 ± 0.5 × 10LJ copies/ng DNA), when compared to survivors with 1.4 ± 0.3 × 10² copies/ng DNA (**Figure 2**). These findings support the hypothesis that shrimp survival following WSSV infection is associated with the host’s ability to reduce viral replication to a tolerable level.

### 3.4. Confirmation of contiguous EVE-468 WSSV fragment in some of the parental broodstock in Crosses 3 and 4

After discovering evidence of linkage of EVE-4, 6 and 8 in Crosses 3 and 4, the question was raised as to whether the 3 EVE were contiguously linked in the parental shrimp as a single EVE fragment (EVE-468) or as separated fragments on the same chromosome. To determine this, PCR tests were carried out using primers designed to amplify a single EVE-468 fragment followed by sequencing of any amplicons that arose from the PCR tests.

#### 3.4.1. Results for presence of a single EVE-468 transcript in Cross 3 (F38 x M17)

**Table 14** for Cross 3 shows results for amplification of a single, linked EVE-468 transcript by PCR from 12 offspring, including 10 surviving and 3 dead shrimp. For the 3 dead offspring, all 3 EVE expressed RNA, but only 1 gave a single EVE-468 PCR product. We concluded that the expressed RNA in these specimens was of positive sense and was thus non-protective. In the 9 remaining shrimp that all survived, 9 gave positive PCR test results for a contiguous EVE-468 fragment. The single exceptional survivor 27f expressed all three EVE-3, -6 and -8 but gave no single EVE-468 amplicon.

**Table 14.**
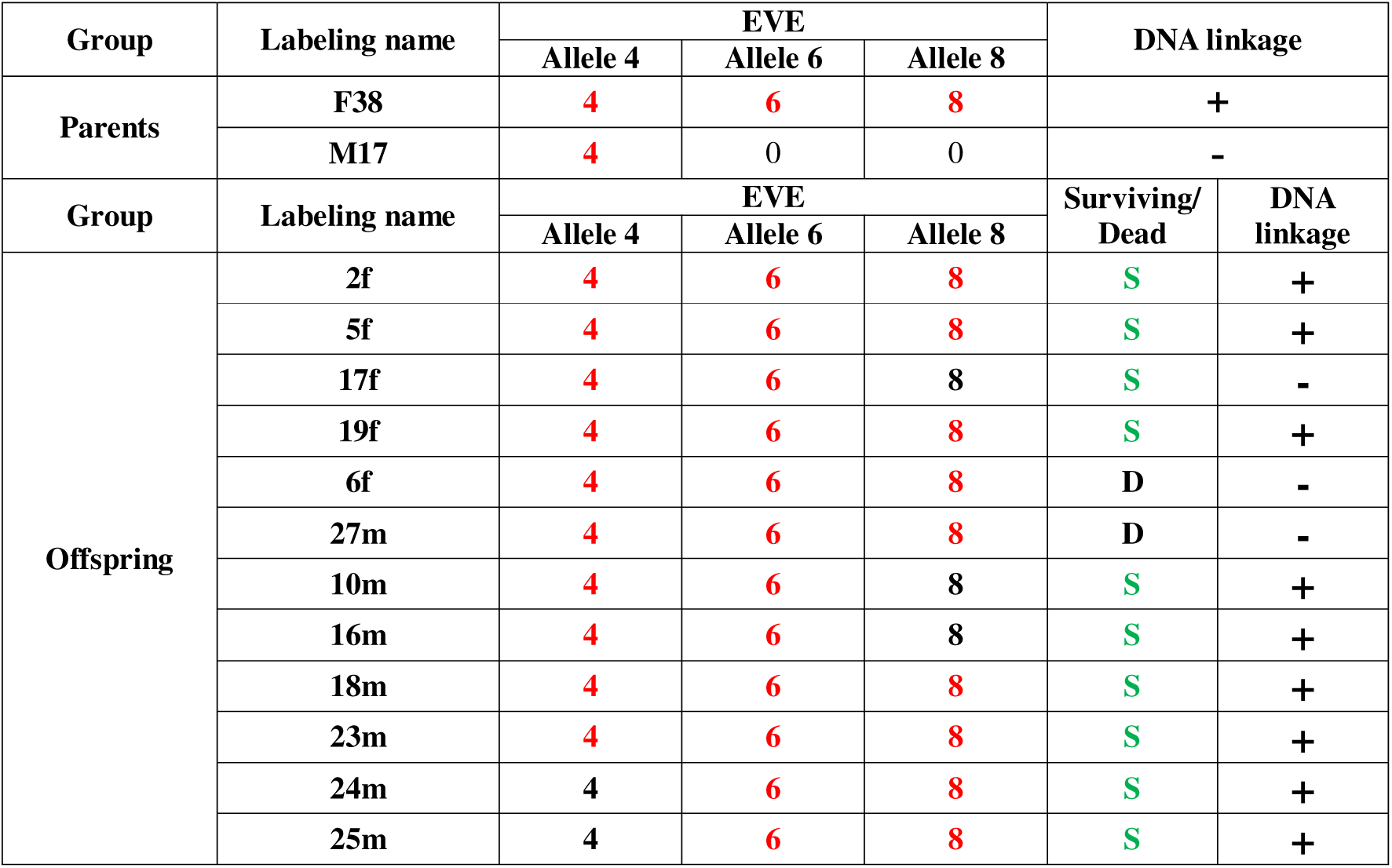
Relationship between linkage of EVE alleles-4,6,8 (EVE-468) in parents and offspring and survival for 13 specimens that could be analyzed from Cross 3 (F38 x M17).

Interpreting these results, it is clear that 9 of the 12 offspring analyzed inherited the linked EVE-468 present in the female parent F38 in the heterozygous state but that RNA expression was not always transmitted together with an allele, indicating some currently unknown and independent control over RNA expression and its sense. Despite this uncertainty, it is clear that 9 of the 10 survivors exhibited this EVE profile.

#### 3.4.2. Results for presence of EVE-468 in Cross 4

**Table 15** for Cross 4 shows results from 13 offspring tested for inheritance of a contiguous WSSV-EVE-468 fragment together with the EVE expression profiles of both parents and offspring. EVE-RNA expressions from both parents occurred as a single genetic element EVE-468 here called an allele in each parent and should give rise to 75% of the offspring positive for expressed EVE-468. The table shows that 11/13 (85%) of the shrimp analyzed were positive for EVE-468. In addition, RNA expression was variable in the offspring, when compared to both parents for which RNA expression was positive. Only 3 of the 13 analyzed shrimp died and 2 of these did not give rise to a contiguous EVE-468 transcript, while the third did, suggesting that it did not produce protective negative sense RNA. Overall, the results indicated that 9/10 = 90% of the survivors in the challenge test had inherited and expressed 2 or 3 of the EVE-4,6,8 alleles. These results confirmed speculations raised in Section 3.5 regarding the possible existence and heritability of a contiguous EVE-468.

**Table 15.** Relationship between linkage of EVE alleles-4,6,8 (EVE-468) and survival in Cross 4 (F64 x M7).

| Group | Labeling name | EVE |  |  | DNA linkage |  |
| --- | --- | --- | --- | --- | --- | --- |
|  |  | Allele 4 | Allele 6 | Allele 8 |  |  |
| Parents | F64 | 4 | 6 | 8 | + |  |
|  | M74 | 4 | 6 | 8 | + |  |
| Group | Labeling name | EVE |  |  | Surviving/Dead | EVE-468 Linked |
|  |  | Allele 4 | Allele 6 | Allele 8 |  |  |
| Offspring | 3f | 4 | 6 | 8 | S | + |
|  | 10f | 4 | 6 | 8 | S | + |
|  | 23f | 4 | 6 | 8 | S | + |
|  | 27f | 4 | 6 | 8 | S | + |
|  | 12f | 4 | 6 | 8 | D | - |
|  | 9m | 4 | 6 | 8 | S | + |
|  | 13m | 4 | 6 | 8 | S | + |
|  | 21m | 4 | 6 | 8 | S | + |
|  | 22m | 4 | 6 | 8 | S | + |
|  | 24m | 4 | 6 | 8 | S | + |
|  | 25m | 4 | 6 | 8 | S | + |
|  | 19m | 4 | 6 | 8 | D | - |
|  | 26m | 4 | 6 | 8 | D | + |

### 3.5. HFRS for infectious WSSV found in pooled cvcDNA of surviving offspring in Cross 4

It was of particular interest for us to determine whether cvcDNA for EVE-4, -6 and -8 could be identified in a pool of the raw-read DNA samples isolated from offspring shrimp prior to their challenge with WSSV. The approach used was based on our earlier work that revealed EVE-4, -6 -8 in *P. vannamei* (Taengchaiyaphum et al. 2024). This allowed us to determine whether comparative analysis of the cvcDNA obtained prior to WSSV challenge of the dead and surviving offspring shrimp from Cross 4 (F64 x M74) would correlate with their EVE profiles. Results from mapping the raw reads with 97-100% DNA sequence identity to an infectious WSSV genome sequence are shown in green outline in **Figure 3**. Also shown in pink outline are cvcDNA fragments that matched ancient WSSV-EVE with no significant nucleic acid sequence identity to extant WSSV. In **Fig. 3A**, cvcDNA fragments from the pooled DNA of dead shrimp show cvcDNA fragments that matched ancient WSSV EVE sequences only and were scattered over the WSSV map (pink outline). The pooled DNA of surviving shrimp **Fig. 3B** gave a different cluster of fragments from non-infectious WSSV sequences (pink outline). However, they also gave a cluster of cvcDNA fragments that matched an extant, infectious WSSV genome (green outline), and fell within the EVE-4,6,8 region previously reported from a WSSV resistant *P. vannamei* stock and copied here in **Fig. 3C**, copied from a previous publication (Taengchaiyaphum et al., 2024) and shown here for easy reference. In summary, no reads matching the genome of infectious WSSV were observed in the dead shrimp group. Although the magnitude of the WSSV fragment hits for the HFRS region in Fig. 3B is lower than in Fig. 3C, their genome locations are almost identical. The results agreed with the PCR results from the crosses and challenges, and they also supported our proposal that these EVE may have been naturally selected in both species via survival from previous WSSV disease outbreaks over time.

**Figure 3.**
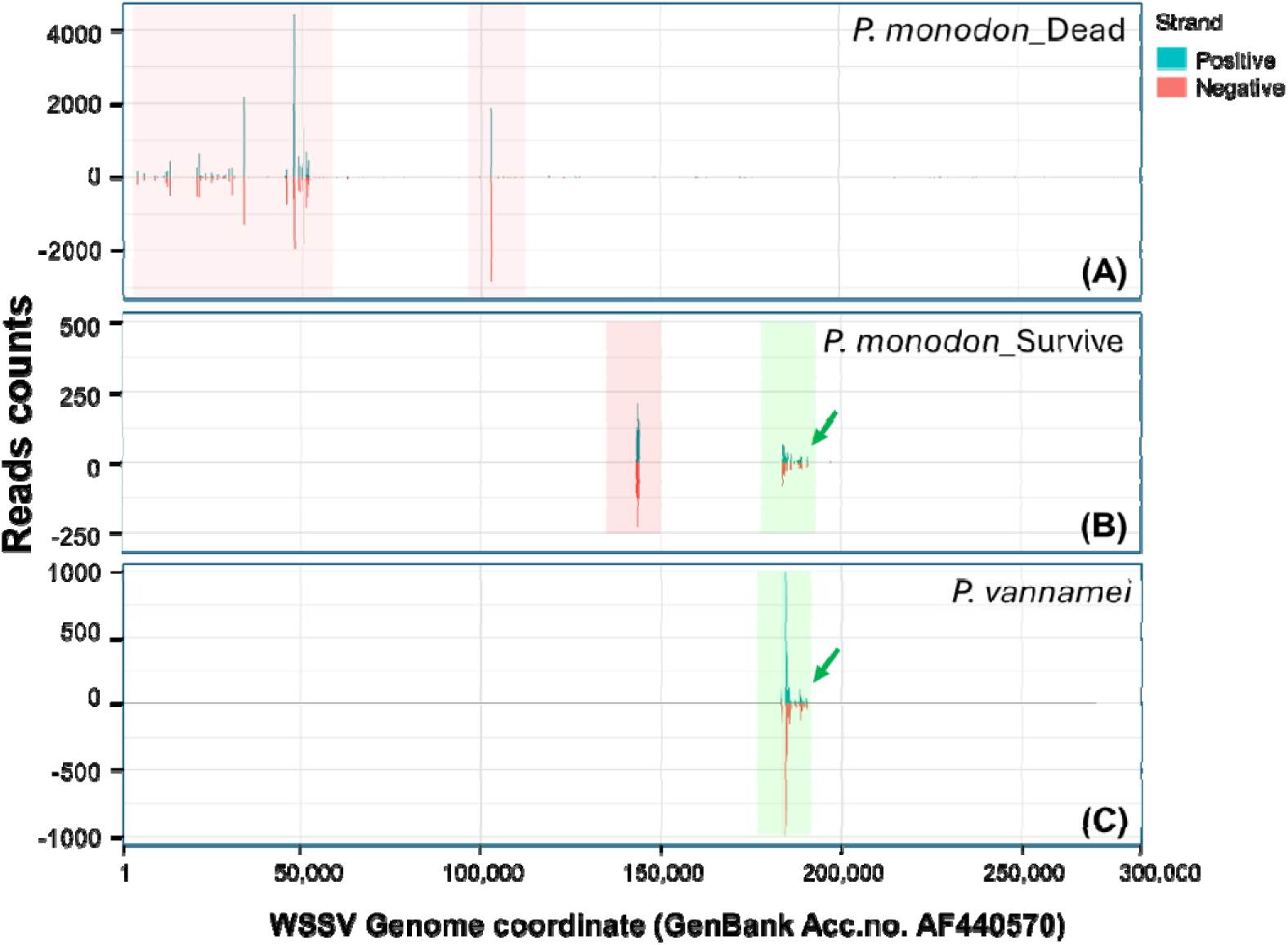
Detection of WSSV-derived circular DNA reads matching infectious WSSV using analysis platform #1 with DNA extracted from WSSV-challenged shrimp prior to challenge. (A) Profile for cvcDNA from the dead shrimp group showing cvcDNA fragments from ancient non-infectious WSSV EVE only (pink outline). (B) Profile from the surviving shrimp showing cvcDNA from both ancient, non-infectious WSSV (pink outline) and from extant, infectious WSSV (green outline). (C) Comparative profile of cvcDNA from a previous study using the same detection approach with circular DNA extracted from a *P. vannamei* breeding stock (Taengchaiyaphum et al., 2024), and showing the same HRFS region as that in *P*. *monodon*. All circular DNA reads for extant WSSV were mapped to the reference genome (GenBank accession no. AF440570).

The absence of infectious WSSV reads in the pooled DNA from dead shrimp from Cross 4 was unexpected, because some individuals did give positive PCR test results for expressed EVE-4,6,8 by PCR analysis (see Table 12). Absence of matching cvcDNA sequences in the dead shrimp group indicated that they did not give rise to cvcDNA. This, in turn, suggested that cvcDNA production itself might be involved in the mechanism for EVE protection. If so, shrimp selected for WSSV tolerance based on PCR tests for HFRS sequences raised the question whether they might also need to be screened to confirm capability to produce cvcDNA from those EVE.

To study this question, we examined 5 surviving and 5 dead shrimp arbitrarily selected from Cross 4 to determine whether there was any relationship between HFRS cvcDNA and survival or death. The results shown in Table 16 reveal that the surviving shrimp pool included 4 shrimp positive for all 3 expressed EVE-4,6,8 while one was positive for all 3 unexpressed alleles.

**Table 16.**
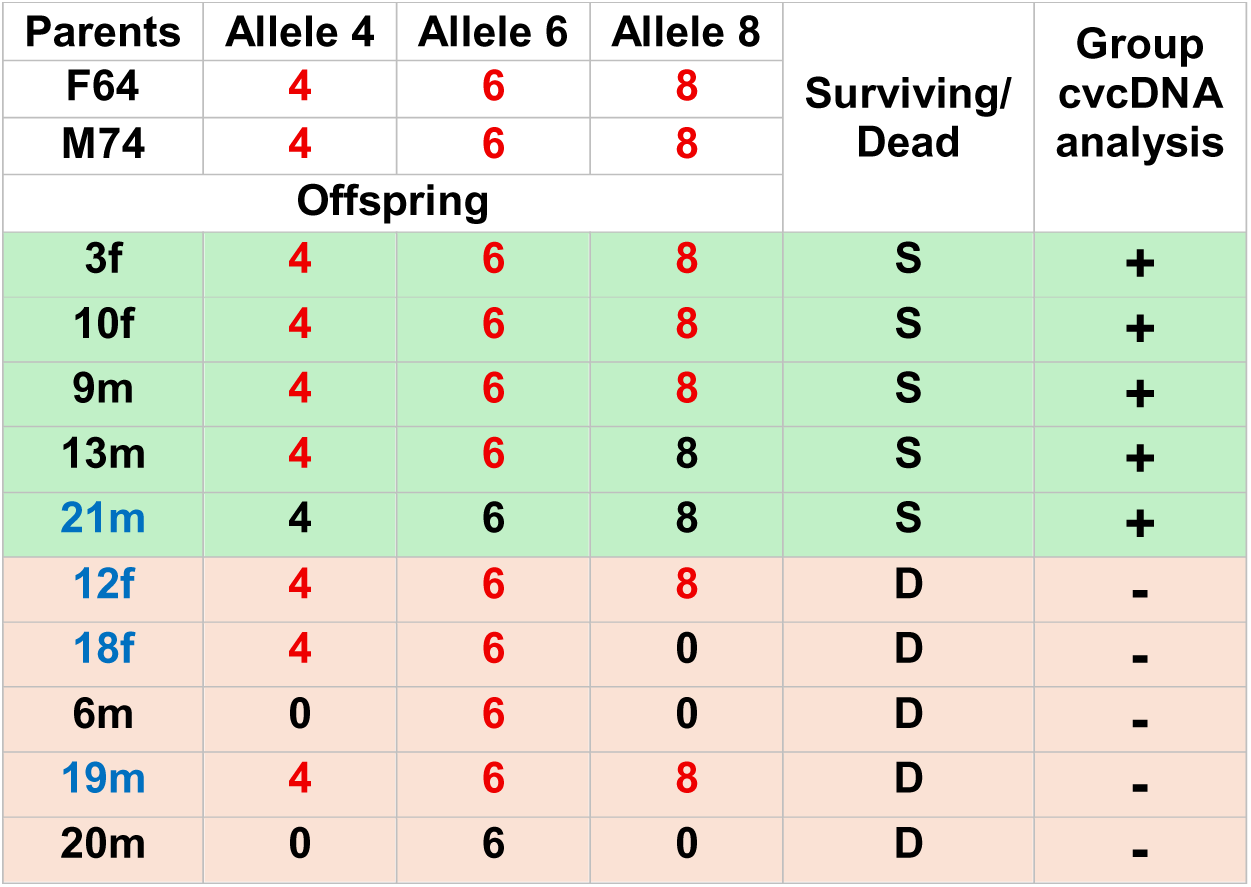
Comparison of cvcDNA EVE-4,6,8 profiles of 5 arbitrarily selected surviving shrimp (green background) and 5 arbitrarily selected dead shrimp (pink background) to compare their group relationship to positive or negative cvcDNA results derived from their pooled DNA preparations.

The dead shrimp included those with mixed EVE-4,6,8 profiles, including 2 shrimp with expression of all 3 alleles, one expressing 2 alleles, another expressing 1 allele and another none. However, this pool of dead shrimp produced no EVE-4.6.8 related cvcDNA. Altogether, these results suggest a key role for cvcDNA entities that match the sequence of genomic EVE in determining the protective capability of those EVE.

## 4. DISCUSSION

### 4.1. Mendelian inheritance of EVE was confirmed

Results from the 4 crosses carried out in this study correspond to those from earlier publications revealing that EVE are inherited in Mendelian fashion (Brock, et al., 2013; Taengchaiyaphum, et al., 2019). This is also supported by whole genome sequencing of *P. monodon* that has revealed scrambled clusters of EVE from the parvovirus IHHNV that occur on more than one chromosome and could thus be transmitted to offspring independently or linked (Huerlimann, et al., 2022) (Taengchaiyaphum, et al., 2022), depending on their genome arrangement and location. This, in turn, reveals that they are susceptible to genetic manipulation in a shrimp breeding program.

### 4.2. WSSV-EVE presence and expression is associated with tolerance to WSSV infection

As previously reported for EVE in a population of *P. monodon* the type of RNA expression (none, negative sense, positive sense or dual positive-negative sense) was variable (Utari, et al., 2017) with the likelihood that only negative sense or dual negative-positive sense RNA expression would result in protection against WSSV. In addition, such EVE were shown to be heritable in Mendelian fashion (Taengchaiyaphum et al, 2019).

In Cross 1 of this limited study, the sense of RNA expression (+/-) was not determined. It was found that EVE-4,5,8 all expressed RNA in the male parent while none were present in the female. All the inherited EVE in its offspring also expressed RNA, matching the male parent from which they originated. Despite at least 1 or 2 being expressed in 9 of 10 offspring, there was no protection against WSSV. This suggested that the transmitted EVE might be non-protective because they did not express negative-sense RNA, which may or may not have been the situation with the parental male that transmitted the 3 EVE. Is it possible that all the RNA expressed in the offspring was of positive sense and thus unable to induce an RNAi response? If any of these EVE were protective for the male parent due to capability for negative-sense RNA expression, that capability was not transmitted to the offspring. If so, it would suggest that the male was capable of transmitting the EVE but unable to transmit the genetic capability for those EVE to produce negative sense RNA. This, in turn, would suggest that control for negative sense expression might reside in mitochondria, since they contain genetic material that males cannot transmit or very rarely transmit to their offspring.

In Cross 2, also with 100% mortality, EVE-4 and -8 were expressed in the male parent while EVE-6 was present without RNA expression. In the offspring, EVE-4 with RNA expression and EVE-6 without were present in one specimen which matched the profile of the male parent. However, EVE-6 and -8 were present in one specimen, both without RNA expression that did not match the male parent in which EVE-8 was expressed. Similarly, presence and RNA expression by EVE-6 in one specimen did not match the parent. These differences in off/on opposites for RNA expression between parental and offspring EVE suggested additional, currently unknown, independent on/off genetic control(s) over EVE RNA expression, and also independent control of sense of RNA expression. Divulging the mechanism(s) for these controls should be an important ongoing research goal. At the same time, searches for shrimp and other crustaceans that carry HFRS-EVE-468 as a contiguous element should be a prime target followed by cvcDNA analysis to ensure that it will be expressed in negative sense before using them for mating experiments.

In contrast, Crosses 3 and 4 revealed that EVE-4,6,8 may sometimes be linked together as a continuous EVE-4, 6, 8 fragment that shows promise as a target for cvcDNA screening of other wild, cultivated or breeding-stocks of crustaceans susceptible to white spot disease. It is also possible that existing breeding stocks might be genetically engineered to incorporate this fragment if it is not already present. However, there is a strong additional need to determine the currently unknown and apparently independent genetic control factors that appear to govern activation and sense of RNA expression from EVE. It is also possible that the cvcDNA isolation protocol of Tassetto et al. 2019 could be used to screen commercial insects such as silkworms and honeybees that have survived previous viral infections but test negative for active infections of the target viruses. It is possible that mapping of DNA raw reads with a high sequence identity to the target virus genome would reveal HFRS regions indicative of naturally selected protective sequences. This might also be possible with organisms from other phyla that do not produce antibodies (Flegel, 2022).

The most startling discovery in this study was the strong association between shrimp survival and their capability to produce cvcDNA of HFRS-EVE. Absence of this capability resulted in death, even with some shrimp specimens that expressed all 3 EVE-4,6,8 alleles. The mechanism behind this phenomenon is unknown to us but appears to be linked to some kind of control over negative-sense RNA expression by the relevant EVE. Understanding the mechanism(s) behind this phenomenon is a prime target for further research.

In summary, there appear to be 2 independent controls related to protective EVE in terms of whether RNA is expressed or not and whether the RNA expression sense will be positive, negative or dual sense. Fortunately, in two of our crosses the three protective EVE were linked and could be amplified as a single PCR product. Thus, it will be a major focus for our further work to select individuals of this nature to simplify the analytical process in experiments on EVE function and expression sense. Given that the HFRS region for WSSV that we discovered has arisen independently in the *P. vannamei* and *P. monodon* stocks used, we recommend that researchers interested in studies on protective EVE for WSSV target this sequence for screening stocks or species for its presence and protective capacity. We believe that the natural occurrence of this sequence region disqualifies it for patenting, so it is open for free general use for all those interested in working with it. In addition, the approach we have used can be applied to detect, screen and test for other potentially protective EVE in other cultured crustaceans and insects.

### 4.3 Proposed, ongoing work

Based on our experience, we recommend that shrimp breeders wishing to explore the possibilities described herein focus on male shrimp for the permanent preservation of key EVE in the form of cryopreserved sperm. In this manner, it should be possible to keep a stock of EVE genetic material that is independent of living shrimp and can be used for artificial insemination. Hopefully this would solve the problem of the relatively short broodstock lifetime in shrimp. Unfortunately, this would not work for any control gene(s) that might occur in mitochondria.

At the end of our study, we had only 1 surviving individual male broodstock shrimp (M225) from Cross 3 to use for ongoing breeding experiments. This individual carried and expressed EVE-468. It was proposed that this male be used to harvest spermatophores for cryopreservation of sperm and for artificial insemination of females lacking the set of EVE alleles described in our earlier publication (Taengchaiyaphum, et al., 2024). This would allow for ongoing experiments to explore the possibility of creating parental stock capable of producing offspring that can be supplied to shrimp farmers as all positive for linked and negatively expressed EVE-4,6,8.

## 5. CONCLUSIONS

This study has revealed that offspring of *P. monodon* carrying WSSV-EVE that express RNA exhibit significantly higher survival when challenged with WSSV than offspring that do not carry WSSV-EVE or offspring that carry such alleles without RNA expression. The WSSV fragments targeted in the study were mapped to a small region of the WSSV genome that was found to yield high frequency raw reads sequences in cvcDNA extracted from WSSV-free *P. vannamei* and *P. monodon* populations that had survived previous exposure to WSSV. These results supported the hypothesis that cvcDNA extracts containing EVE raw-read fragments that cluster in HFRS regions may serve as indicators of natural, genetically selected EVE that are protective against white spot disease. This approach may be useful to search shrimp breeding stocks for EVE protective against not only WSSV but also other shrimp viruses. Key targets of the ongoing work must be to understand the currently unknown independent controls over EVE-RNA expression or not and over negative sense or dual sense RNA transcripts or not, and how these might be manipulated to produce shrimp postlarvae all of which would be naturally tolerant to WSSV.

## Data Declaration

The datasets used and/or analyzed during the current study are available from the corresponding author on reasonable request.

## Acknowledgement

This research was funded by the National Research Council of Thailand (NRCT) through the High-Potential Research Team Grant Program (Grant No. N42A650869) and by the Mahidol University (Fundamental Fund: fiscal year 2025 by National Science Research and Innovation Fund (NSRF)) (Grant no. FF-115/2568).

